# Full-length 16S ribosomal RNA gene sequencing reveals dynamics of tick-adapted and environmentally derived bacteria in the microbiome of the black-legged tick, *Ixodes scapularis* in Nova Scotia, Canada

**DOI:** 10.64898/2026.06.11.731624

**Authors:** Saffi Sangster, Katherine A. Dunn, Emma Phelan, Jessica Latimer, James Kho, Tatiana Rossolimo, Amal E. Nabbout, Shelley A. Adamo, John M. Archibald

## Abstract

Lyme disease is a tick-borne illness caused by the spirochaete bacterium *Borrelia* (*Borreliella*) *burgdorferi*. The black-legged tick *Ixodes scapularis*, which transmits *B. burgdorferi* and several other human pathogens, is endemic to the eastern United States and, due to climate change, is rapidly expanding into central and eastern Canada. Amplification and sequencing of bacterial DNA from *I. scapularis* is increasingly used to monitor the presence and abundance of *B. burgdorferi* and associated bacteria. However, variation in the nature of molecular data collected across studies presents challenges for analysis and interpretation. Here we use full-length Oxford Nanopore 16S ribosomal RNA gene amplicon sequencing to characterize the microbiome of *I. scapularis*, with an explicit focus on distinguishing between tick-adapted bacteria (endosymbionts and pathogens) and environmentally acquired bacteria (external sources, including soil, vegetation or vertebrate hosts). We show that environmental dominance strength differs between these two ecological classes of bacteria, and that environmental dominance does not appear to represent stochastic background alone; environmentally derived bacterial taxa detected in tick microbiomes are not mere contaminants. Paired soil microbiome profiling from tick collection sites will be required to test whether environmental dominance and associated co-occurrence structure track with seasonal changes in exposure and environmental microbial populations.

## Introduction

The black-legged tick, *Ixodes scapularis*, is the primary vector of several bacterial pathogens of major public health importance in North America. Most notably, it transmits *Borrelia* (*Borreliella*) *burgdorferi*, the causative agent of Lyme disease, which is the most reported vector-borne disease in the United States and Canada (CDC, 2024; Steere *et al*. 2016). In addition to *B. burgdorferi, I. scapularis* can also carry other clinically significant bacterial pathogens, including *Anaplasma phagocytophilum* and *Ehrlichia chaffeensis*, as well as protozoan parasites such as *Babesia microti* (Eisen & Eisen 2018; Paddock & Childs 2003). The incidence of these tick-borne diseases and the geographic range of the ticks carrying them have increased substantially over recent decades. In Nova Scotia, Canada, for example, public health surveillance data reveal marked increases in the detection of tick-borne ‘notifiable diseases’ between 2019 and 2024 alone (Nova Scotia Health, 2024). These observations underscore the need to better understand biological factors within ticks that may influence pathogen acquisition, persistence, and transmission (Eisen & Eisen 2018; Ogden *et al*. 2014).

Beyond their role as pathogen vectors, *I. scapularis harbors* complex bacterial assemblages derived from multiple ecological sources, including vertically transmitted endosymbionts, horizontally acquired tick-adapted pathogens, and bacteria acquired from the surrounding environment through contact with soil, vegetation, and vertebrate hosts (Narasimhan & Fikrig 2015; Bonnet *et al*. 2017; Tawidian *et al*. 2025). Endosymbionts such as *Rickettsia* are intracellular, transmitted from parent to offspring, and often dominate the tick microbiome, where they are considered stable and persistent community members (Sassera *et al*. 2006; Kurtti *et al*. 2015).

Tick-borne pathogens, including *Borrelia, Anaplasma*, and *Ehrlichia*, are also tick-adapted but differ in transmission and ecological role. These bacteria are horizontally acquired during blood feeding, replicate within tick tissues as part of a vertebrate—tick lifecycle, and are not free-living environmental organisms (Dumler *et al*. 2001 ; Radolf *et al*. 2012). Together, endosymbionts and tick-borne pathogens represent biologically integrated components of the *Ixodes* microbiome, despite their distinct functional roles.

In contrast, numerous DNA sequencing-based studies have reported frequent detection of soil- and plant-associated taxa (e.g., *Pseudomonas, Bacillus*) as well as vertebrate host-associated bacteria (e.g., *Streptococcus, Staphylococcus*) in the Ixodes microbiome (Zolnik *et al*. 2016; 2018; Thapa *et al*. 2019a and b). The presence of these taxa does not necessarily imply persistence or ecological integration within the tick, and dominance by environmentally acquired bacteria—here defined as bacteria derived from external sources, including soil, vegetation, and vertebrate hosts—is often interpreted to be a consequence of transient exposure rather than stable colonization (Ross *et al*. 2018; Lejal *et al*. 2021). However, it remains unclear whether these environmentally acquired taxa occur primarily as stochastic ‘background’ or whether they form coordinated assemblages that represent distinct microbiome states within ticks. Such coordinated environmental assemblages, if present, may reflect ecological processes linked to tick behavior and exposure, including seasonal variation in questing activity and environmental contact, suggesting a potential role for time of year in shaping microbiome dominance states.

Interpretation of tick microbiome composition is further complicated by variation in the type of molecular sequence data collected and analyzed. PCR amplification and sequencing of the 16S ribosomal RNA (16S rRNA) gene using so-called ‘short-read’ technologies such as Illumina typically target only small variable regions of the gene. By nature, short amplicons contain limited phylogenetic signal and thus can result in the collapse of otherwise distinct bacteria taxa into groupings with shared labels, thus obscuring closely related species, and limiting resolution among ecologically divergent groups, particularly within genera that are common in tick microbiomes (e.g., *Pseudomonas, Rickettsia, Borrelia*). Full-length 16S sequencing using long-read DNA sequencing platforms such as Oxford Nanopore has the potential to improve taxonomic resolution and provide more consistent genus- and species-level assignments, thus facilitating more accurate assessment of dominance structure and co-occurrence patterns across samples.

In this study, we use full-length nanopore 16S amplicon sequencing to characterize dominance structure and community organization in the microbiome of *I. scapularis* in Nova Scotia, with explicit distinction between tick-adapted bacteria (endosymbionts and pathogens) and environmentally acquired taxa. We examine whether dominance strength differs between these ecological classes, assess how dominance state relates to co-occurrence structure, and evaluate how *Borrelia*-associated relationships vary across microbiome states. By focusing on dominance rather than prevalence alone, this work aims to clarify whether environmentally acquired bacteria are primarily stochastic background or represent coordinated assemblages with potential ecological and indirect biomedical relevance.

## Methods

### Tick collection and DNA extraction

Adult *Ixodes scapularis* ticks were collected in May 2024 in a field located in Burnside, Dartmouth, Nova Scotia, by drag sampling using a white cloth. Ticks were identified as adult males (n = 2) or adult females (n = 5), placed in 50 mL Falcon tubes with moistened cotton to prevent desiccation, and stored at 4°C.

Immediately before extraction, ticks were washed in ethanol to remove surface contaminants and air-dried for approximately 5 min in a sterile Petri dish. Ticks were bisected using ethanol-sterilized tweezers and a razor blade, and tissue fragments were incubated in 180 μL ATL buffer with 20 μL Proteinase K at 56°C with shaking (500 rpm) overnight (∼16 h) following established protocols (Ammazzalorso *et al*. 2015). Genomic DNA was extracted using the DNeasy Blood and Tissue Kit (Qiagen) according to the manufacturer’s instructions. DNA quality was assessed using a NanoDrop spectrophotometer, and DNA concentration was quantified using a Qubit fluorometer.

### Additional tick samples and DNA handling

DNA from an additional 81 adult female *I. scapularis* ticks was obtained from previously published (Nabbout *et al*. 2023) and ongoing studies conducted in the Adamo laboratory (Dalhousie University). Fifty-one ticks were collected in October 2020 from Burnside, Dartmouth, Nova Scotia. Fifteen ticks were used as part of a thermopreferendum experiment (10-50°C; unpublished), and 36 were subjected to controlled temperature incubations over winter and retrieved in March 2021 .

The remaining ticks were collected in October 2020 from Thomas Raddall Provincial Park (Port L’Hebert, Nova Scotia) and were used in an overwintering study, with microcosms placed at the Harrison Lewis Centre until retrieval in March 2021 (Nabbout *et al*. 2023). Following retrieval, all ticks were frozen whole at −80°C until DNA extraction in 2022. For the 81 Adamo lab ticks no pre-extraction ethanol wash was performed.

All 81 samples were previously screened for the presence of *Borrelia burgdorferi* by qPCR as described previously (Adamo *et al*. 2022). Remaining DNA was stored at −80°C until use in the present study.

### DNA storage quality control

To assess DNA stability following long-term storage of Adamo lab ticks, five representative samples spanning qPCR-positive, -negative, and indeterminate outcomes were re-quantified in 2024 using a Qubit fluorometer. Re-quantified concentrations closely matched original measurements prior to freezing. PCR amplification was performed and confirmed by agarose gel electrophoresis. DNA concentrations for all tick samples are shown in Table S1 .

### DNA Sequencing

DNA was amplified using the Oxford Nanopore Technologies (ONT) 16S Amplicon Barcoding Kit (SQK-16S114.24) with universal primers 27F and 1492R, which include 5’ tags for attachment of rapid sequencing adapters. Samples were multiplexed in pools of 15-20 using barcoded primers supplied with the kit. Sequencing libraries (50 fmol) were loaded onto R10.4.1 flow cells (FLO-MIN114) and sequenced on an ONT MinION device for up to 72 h. When sufficient library material remained, additional library was loaded at the end of the initial run and sequencing was extended for up to an additional 72 h.

Basecalling was performed using Dorado v0.7.2 (Oxford Nanopore Technologies) with the super-accuracy model (sup@v5.0.0). Demultiplexing and adapter trimming were carried out using Porechop v0.2.4 (Wick *et al*. 2017).

### Processing nanopore metabarcoding data

Nanopore 16S rDNA reads were processed using PRONAME (Dubois *et al*. 2024). Reads were imported using proname_import and filtered to retain sequences between 1,300 and 1,600 bp in length with a minimum Phred quality score of 20 (proname_filter). To reduce sequencing error, reads were clustered at 97% sequence identity using MMseqs2 (Steinegger & Söding 2017). For each cluster, a centroid sequence was refined by polishing with up to 300 randomly selected subreads using Medaka (github.com/nanoporetech/medaka) (proname_refine).

Chimeric sequences were removed, and taxonomic assignment was performed using QIIME2 (Bolyen *et al*. 2019) against a curated reference database of complete bacterial 16S rDNA sequences from NCBI (accessed Aug 1, 2025), supplemented with additional sequences to support species-level identification. The database was formatted using RESCRIPt (Robeson *et al*. 2021), and reference sequences shorter than 1,200 bp were excluded. Resulting abundance tables were used for downstream analysis.

### Microbiome data filtering, normalization, and dominance analysis

Amplicon sequence abundance tables generated from full-length 16S sequencing were analyzed using R (version 4.4.1). Initial analyses were conducted using read count tables at the genus level. To reduce the influence of extremely low-abundance observations potentially caused by sequencing noise, counts representing less than 0.01% of the total reads within a sample were set to zero. In addition, taxa present in fewer than three samples were excluded from downstream analyses.

Dominance structure was quantified on a per-sample basis, defined as the relative abundance of the most abundant taxon within a sample. Dominant taxa were classified according to ecological category as either tick-adapted (including endosymbionts and tick-borne pathogens) or environmentally acquired. Dominance strength was further categorized as *strong* or *weak* based on the margin between the most abundant and second most abundant taxa, with strong dominance defined as a dominance margin exceeding 0.5. To assess the sensitivity of the results to the definition of dominance, analyses were repeated using an alternative dominance margin threshold of 0.3.

### Alpha diversity analyses

Alpha diversity metrics, including Shannon diversity and Pielou’s evenness, were calculated using count data following low-abundance filtering but prior to prevalence-based taxon removal, as removal of rare taxa can bias diversity estimates. Diversity calculations were performed using the vegan package in R (Oksanen *et al*. 2026). Comparisons between dominance strength categories (strong vs weak) and ecological source categories (tick-adapted vs environmental) were preformed using nonparametric tests (α = 0.05) due to non-normal distributions. For two-group comparisons, Wilcoxon rank-sum tests were used. Where four dominance-source categories were compared (env-strong, env-weak, tick-strong, tick-weak), the Kruskal-Wallis test was performed followed by pairwise Wilcoxon rank-sum tests with false discovery rate correction for multiple comparisons.

### Beta diversity analyses

Beta diversity was assessed using Bray-Curtis dissimilarity (based on relative abundance). Dissimilarity matrices were computed using the vegan package in R. Community differences among groups were tested (α = 0.05) using permutational multivariate analysis of variance (PERMANOVA; implemented with adonis2 function, 999 permutations). Models were run separately for dominance strength (strong vs weak), ecological source (tick-adapted vs environmental) and combined four level dominance-source grouping. To assess homogeneity of multivariate dispersion, PERMDISP (betadisper, 999 permutations) analyses were conducted. Principle coordinate analysis (PCOA) was used to visualize Bray-Curtis dissimilarities among samples.

### Co-occurrence and association analyses

To evaluate patterns of taxon co-occurrence, compositional effects were addressed using a centered log-ratio (CLR) transformation. Prior to transformation, a pseudocount of one was added to count data to accommodate zero values. Within group correlation matrices were calculated using Pearson correlations on CLR-transformed data. Networks were constructed by retaining taxon-taxon correlations exceeding an absolute threshold of |r| ≥ 0.3. To reduce instability in correlation estimates, taxa were required to be present in ≥ 20% of samples within each group prior to network construction. Network analyses were visualized and summarized using the igraph package (v1.2.6; Csárdi & Nepusz 2006). Network properties, including node count, edge count, density, and average degree, were calculated for descriptive comparison across dominance-ecological states.

To assess associations between *Borrelia* and other taxa, correlations between *Borrelia* and all other taxa were extracted from group-specific correlation matrices. Taxa were retained if they showed a correlation magnitude of at least r≥0.3 in at least one dominance state. For sensitivity analysis, networks were reconstructed after excluding *Borrelia*-dominant samples to evaluate whether correlation structure was driven by dominance events. Because correlation-based network inference does not imply causation and can be sensitive to sample size, network metrics were interpreted descriptively rather than subjected to formal statistical comparisons.

Among the 88 *I. scapularis* tick samples only two were from male ticks, which meant comparisons of tick microbiomes based on sex could not be examined.

## Results

### DNA sequencing output and taxonomic resolution

DNAs from a total of 88 *I. scapularis* ticks collected in Nova Scotia were used as substrate for PCR amplification of full-length 16S rRNA genes and sequencing with Oxford Nanopore technology. Amplicons were ∼1,450 nucleotides in length, a substantial increase over the Illumina-based amplicons typically used to detect the presence and abundance of *Borrelia* and other pathogens in ticks relative to other bacterial taxa. Total reads per sample ranged from 15,538 to 2,186,246 with a median of 480,819. After quality control, samples contained a median of 344,120 reads (range 9430-1,560,753) (Table S2). After filtering (removal of taxa present in <0.01% of reads per sample and in <3 samples) the dataset contained 153 taxa (genus-level), with a median of 25 taxa per sample (range 3 to 73; Table S2).

### *Ixodes scapularis* microbiome composition

Across tick samples, bacterial taxa varied substantially in both prevalence and mean relative abundance (Fig. 1A). The genus *Rickettsia* exhibited the highest mean relative abundance (58%) but was the third most prevalent taxon (89.8%). In contrast, *Pseudomonas* and *Caballeronia* were detected in a greater proportion of samples (92.0% and 90.9%, respectively) yet contributed substantially lower mean relative abundances (14.6% and 5.3%). Several additional environmentally associated genera, including *Luteibacter, Sphingomonas*, and *Methylobacterium*, were detected in more than 77% of individual ticks but each represented less than 3.5% mean relative abundance (Fig. 1A). Thus, many taxa were widespread but typically present at a low proportional abundance.

**Figure 1.**
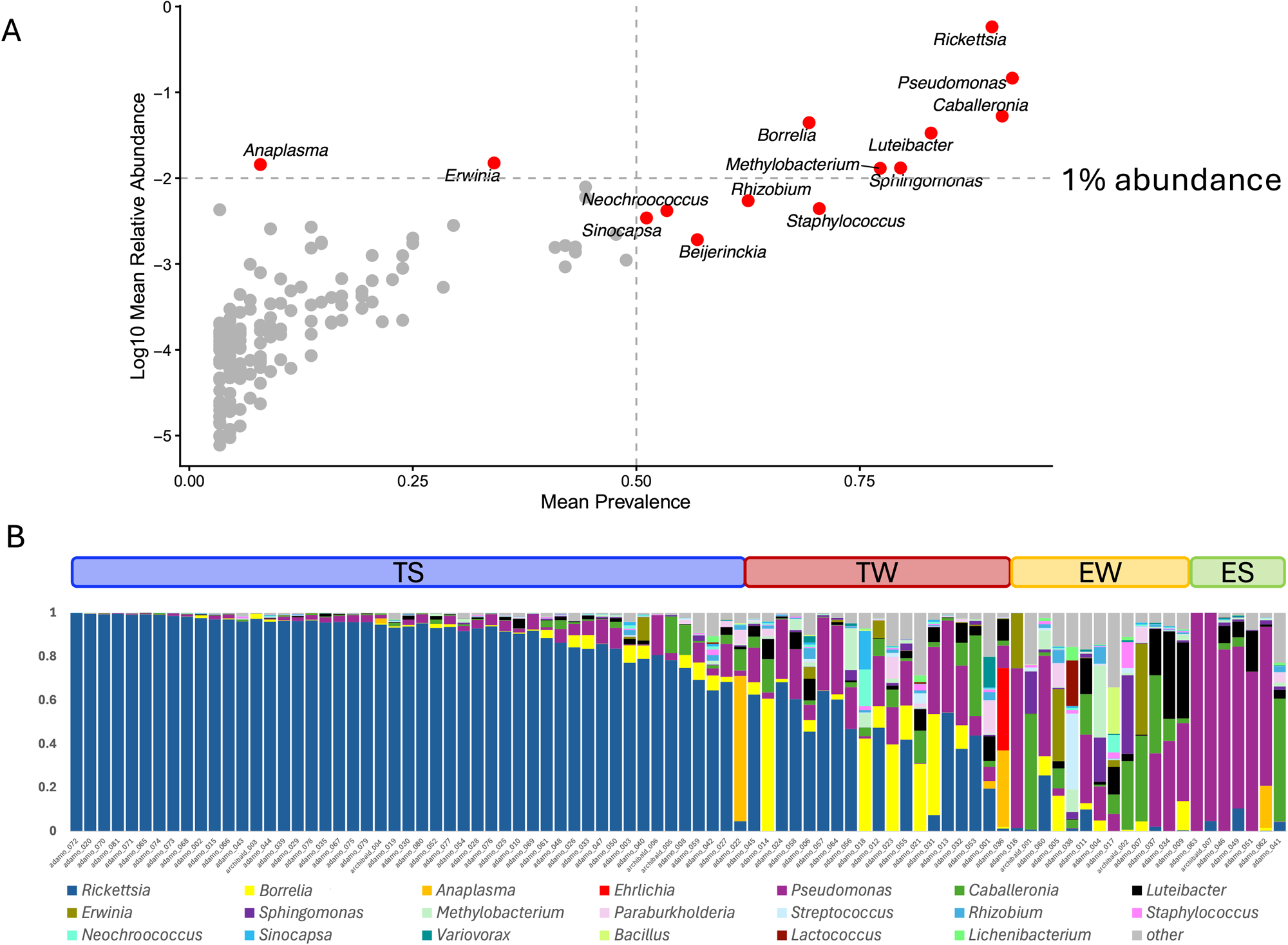
Composition and prevalence of microbial community members in the tick *Ixodes scapularis* in Nova Scotia, Canada. **(A)** Mean prevalence vs mean relative abundance (log10) of tick samples. Taxa present with mean abundance >50% or mean relative abundance >1 % are labeled and colored in red. **(B)** Barplot of relative abundance for each sample. Samples are ordered by ecological class and dominance strength. TS =tick adapted-strong dominance; TW = tick-adapted-weak dominance; EW=environmental-weak dominance; ES=environmental-strong dominance.

The tick-borne pathogen *Borrelia* was detected in 69% of samples, with a mean relative abundance of 4%. *Anaplasma* (7/88 samples) and *Ehrlichia* (3/88 samples) were detected at lower prevalence, 7.95 and 3.4% respectively. Comparison of these sequencing-based data to previously published qPCR screening of a sub-set of the samples analyzed herein (Nabbout *et al*. 2023) showed strong concordance. Among 53 samples qPCR-positive for *Borrelia*, 50 (94.3%) yielded actual *Borrelia* sequences in our study, whereas 18 of 22 qPCR-negative samples (81.8%) lacked detectable *Borrelia* reads (Table S2). The small number of discordant cases likely reflects low-abundance infections near assay detection limits rather than systematic disagreement between methods.

### Taxonomic dominance structure varies across tick samples

Dominance was quantified using the Berger-Parker index (Table S3), which identifies the most abundant taxon within each sample. This index was used solely to define dominant taxa and dominance strength, rather than as a general diversity metric. Dominance strength was calculated as the difference in relative abundance between the most abundant and second most abundant taxa, with samples classified as strongly dominated (?0.5) or weakly dominated (<0.5) (Table S3). Eleven different taxa were found to be dominant in at least one tick with Rickettsia dominant most often followed by *Pseudomonas*, and *Borrelia* (Table 1). Dominant taxa were classified according to ecological category as either tick-adapted (including endosymbionts and tick-borne pathogens) or environmentally acquired, as conceptualized by others (Couper *et al*. 2019; Narasimhan *et al*. 2021 ; Maldonado-Ruiz 2024). At a margin of 0.5, 68 samples were dominated by tick-adapted taxa (49 strong, 19 weak) while 20 samples were dominated by environment-derived taxa (7 strong, 13 weak) Fig 1 B, Table S3.

**Table 1.** Microbiome of *Ixodes scapularis*. Table shows identified dominant taxa with number of samples each taxon was dominant in, along with mean (min, max) relative abundance of that taxon in those samples, and mean marginal differences in those samples.

| **Taxa** | **Number of samples dominant in (strong/weak)** | **Mean relative abundance when dominant (min, max)** | **Mean marginal difference (min, max)** |
| --- | --- | --- | --- |
| **Tick-adapted** |  |  |  |
| *Anaplasma* | 1 (1/0) | 0.6651 (na<sup>1</sup>) | 0.5673 (na) |
| *Borrelia* | 5 (0/5) | 0.4400 (0.31, 0.61) | 0.2491 (0.16, 0.45) |
| *Ehrlichia* | 1 (0/1) | 0.3764 (na) | 0.0261 (na) |
| *Rickettsia* | 61 (48/13) | 0.8227 (0.196, 0.999) | 0.7342 (0.04, 0.999) |
| **Environmental** |  |  |  |
| *Bacillus* | 1 (0/1) | 0.2125 (na) | 0.0857 (na) |
| *Caballeronia* | 3 (1/2) | 0.4828 (0.36, 0.56) | 0.2909 (0.02, 0.51) |
| *Erwinia* | 2 (0/2) | 0.3732 (0.33, 0.42) | 0.0954 (0.02, 0.17) |
| *Methylobacterium* | 1 (0/1) | 0.3301 (na) | 0.1275 (na) |
| *Pseudomonas* | 11 (6/5) | 0.6597 (0.31, 0.999) | 0.4730 (0.009, 0.998) |
| *Sphingomonas* | 1 (0/1) | 0.3633 (na) | 0.0450 (na) |
| *Streptococcus* | 1 (0/1) | 0.3523 (na) | 0.1403 (na) |

<sup>1</sup> na - not applicable, only one sample so minimum and maximum are not shown.

To assess sensitivity to the definition of dominance, analyses were repeated using a lower dominance margin threshold of 0.3. This resulted in a shift in strong tick-adapted taxa from a total of 49 to 55 samples, and strong environment-derived from 7 to 9.

### Alpha diversity differs by dominance state

Shannon diversity across samples ranged from 0.003 to 2.29 (mean = 0.94). As expected, this diversity measure differed significantly between strongly and weakly dominated samples (Fig. 2A; χ2 = 48.516, df= 1, p = 3.28 × 10−2), reflecting reduced evenness in samples dominated by a single bacterial taxon. Consistent with this interpretation, Pielou’s evenness was significantly lower in strongly dominated samples than in weakly dominated ones (Fig. 2B; χ2 = 50.469, df= 1, p = 1 .21 × 10−12), indicating that dominance strength is the primary driver of alpha diversity patterns.

**Figure 2.**
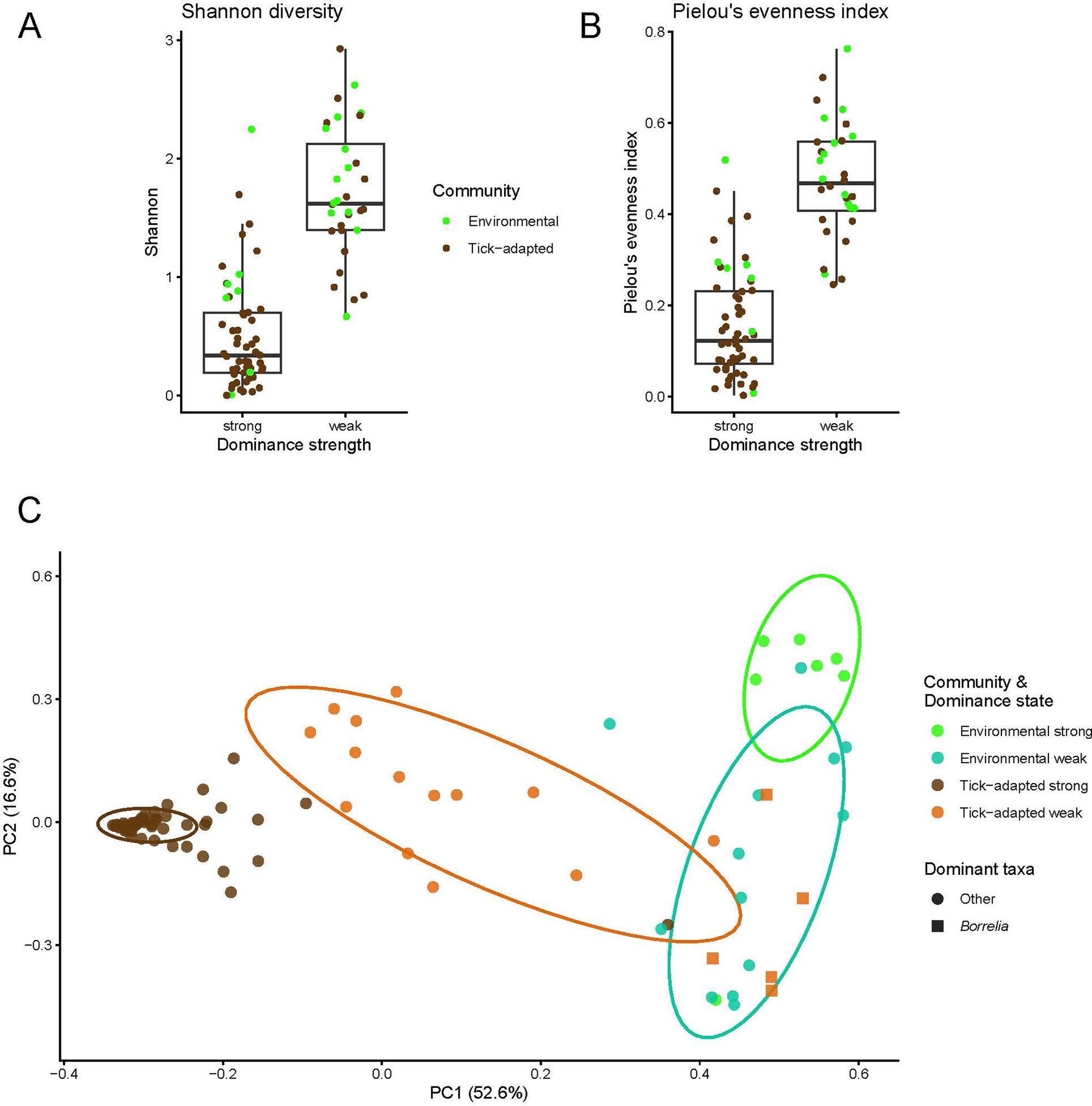
Microbiome diversity in Ixodes scapularis. (**A)** Alpha diversity plot of Shannon diversity by dominance structure (χ2 = 48.516, df= 1, p = 3.28 × 10−12) **(B)** Alpha diversity plot of Pielou’s evenness by dominance structure (χ2 = 50.469, df= 1, p = 1 .21 × 10−12). Dominance was defined by a marginal difference between top two most abundant taxa with strong ≥ 0.5 and weak dominance <0.5. Colors of dots represent ecological category of top taxa with green being environmentally acquired taxa and brown as tick-adapted taxa. (C) Bray-Curtis beta diversity plot showing samples colored by ecological category and dominance structure (Permanova R2= 52.4, p = 0.001 ; PC1 and PC2 explained 52.6% and 16.6% of the variance). Lime green = environmental-strong dominance; teal green=environmental-weak dominance; brown=tick-adapted-strong dominance; orange=tick-adapted weak dominance. Samples where Borrelia was the dominant taxa are indicated as squares.

When tick samples were grouped solely by the ecological origin of the dominant bacterial taxon, alpha diversity also differed significantly between tick-adapted and environmentally dominated microbiomes (Fig. S1). However, stratification by dominance strength revealed that this effect was not uniform across dominance states (Fig. S2). Significant differences were only detected between different dominance strengths of origins, with the exception of Pielou’s evenness, which was significant between origins among strongly dominated samples, but this was weaker than the effect of dominance strength itself (Table S4). These patterns were robust to a more permissive dominance definition (margin>0.3; see Table S4).

### Beta diversity reveals dominance-associated microbiome community structure

Bray—Curtis ordination revealed clear, statistically significant differences in tick microbiome community structure associated with both dominance strength and the ecological origin of the dominant taxon (Fig. 2C). PERMANOVA indicated that dominance strength alone explained 23.7% of community variation (p = 0.001 ; Fig S3A), while ecological source explained 32.8% (p = 0.001 ; Fig S3B). When dominance strength and ecological source were considered jointly, 52.4% of the variation in community structure was explained (p = 0.001) (Fig. 2C).

Tests for homogeneity of dispersion were significant for all groupings (PERMDISP, p < 0.01), indicating that differences among dominance states reflect both shifts in microbiome community composition and differences in within-group heterogeneity. Strongly dominated samples formed more tightly clustered groups, whereas weakly dominated and environmentally dominated samples exhibited greater dispersion in ordination space. Notably, tick samples in which *Borrelia* was the dominant taxon tended to cluster with environmentally dominated samples in ordination space, contributing to increased dispersion within the tick-adapted group.

### Co-occurrence network structure reflects dominance intensity and pathogen embedding

To evaluate coordinated variation among taxa, co-occurrence networks were constructed within each dominance group using CLR-transformed counts and Pearson correlations (|r| ≥ 0.3), retaining taxa present in ≥20% of samples within each group. Network connectivity differed strongly by dominance state. Tick-strong samples, i.e., ticks with microbiomes dominated by tick-adapted bacteria, yielded sparse networks (25 nodes, 41 edges; density = 0.14; mean degree = 3.3; Fig 3A), whereas tick-weak samples produced substantially more connected networks (55 nodes, 442 edges; density = 0.30; mean degree = 16.1 ; Fig 3B), comparable to environmentally dominated samples (50 nodes, 367 edges; density = 0.30; mean degree = 14.7; Fig 3C). Thus, dominance intensity was the primary determinant of network connectivity.

**Figure 3.**
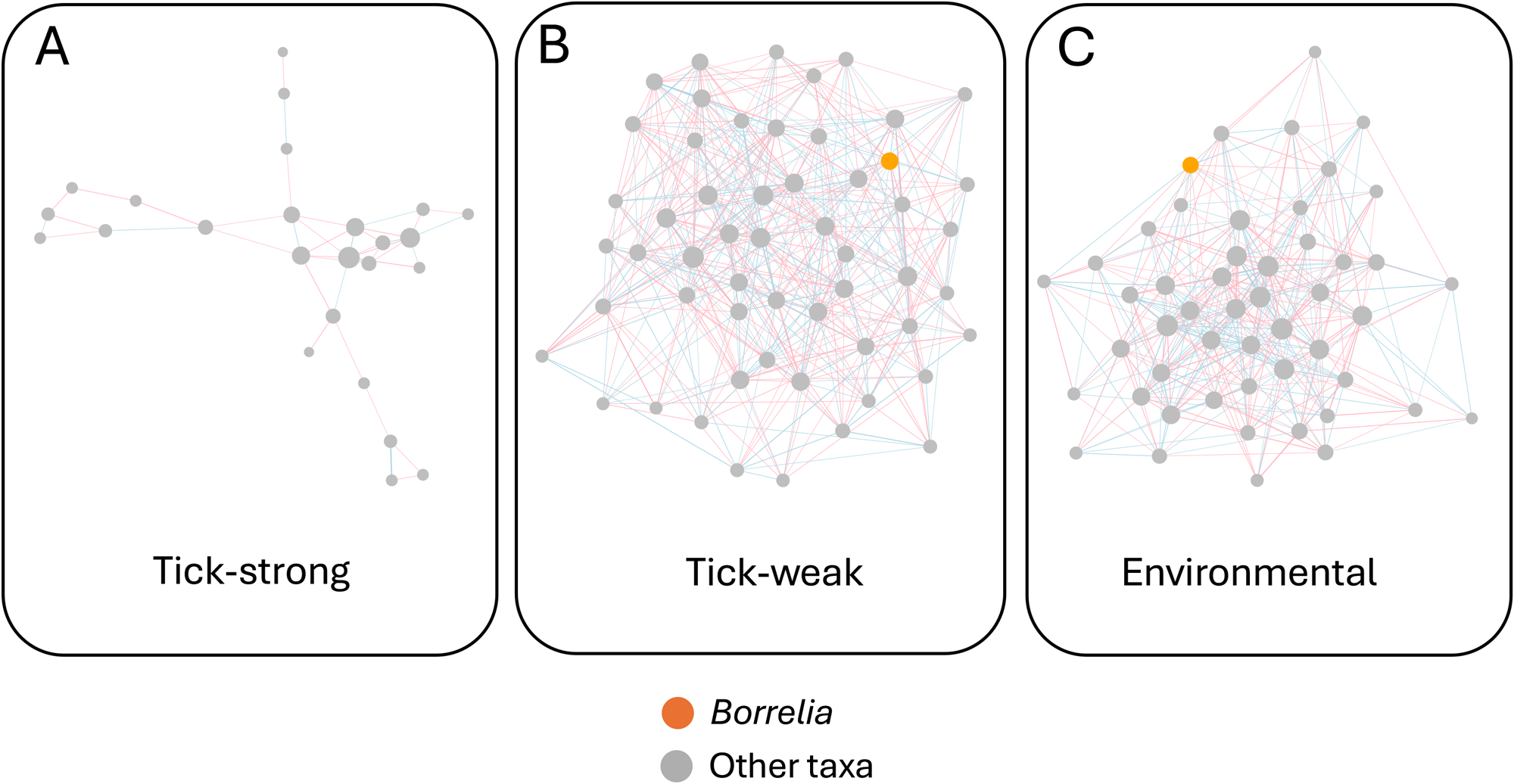
Co-occurrence network structure differs among dominance states in the microbiome of *Ixodes scapularis*. Samples were grouped by dominance state (tick-strong, tick-weak, environmental). Within each group, taxa present in ≥20% of samples were retained, count data were centered log-ratio (CLR) transformed, and Pearson correlations were calculated. Edges were defined using a correlation threshold of |r| ≥0.3. **(A)** Co-occurrence networks for tick samples that were strongly dominated by tick-adapted taxa (margin >0.5; 25 nodes, 41 edges; density = 0.14; mean degree = 3.3). **(B)** Co-occurrence networks for tick samples that were weakly dominated by tick-adapted taxa (margin <0.5; 55 nodes, 442 edges; density = 0.30; mean degree = 16.1). **(C)** Co-occurrence networks for tick samples that were dominated by environmental taxa (50 nodes, 367 edges; density = 0.30; mean degree = 14.7). Node size is proportional to within-network degree (normalized independently for each network). Node corresponding to Borrelia are colored orange with all other nodes colored grey. Edges corresponding to positive correlations are shown in blue, and negative correlations in red.

To examine pathogen embedding within networks, we extracted the *Borrelia* correlation profile (i.e., the *Borrelia* row of the group-wise correlation matrices) from each group-specific matrix. In tick-strong samples, Borrelia showed no associations exceeding |r| ≥ 0.3, consistent with reduced coordinated variation in strongly dominated communities. In contrast, in tick-weak and environmentally dominated samples, *Borrelia* exhibited multiple positive and negative correlations with subsets of taxa (Fig. 3B and C, blue and red lines, respectively). In environmentally dominated samples, *Borrelia* was negatively correlated with several soil-associated genera including *Variovorax, Caballeronia*, and *Luteibacter*, and positively correlated with taxa such as *Erwinia* and *Legionella* (Table S5). In tick-weak samples, a similar pattern of structured covariance was observed: *Borrelia* showed negative correlations with *Anaerococcus* and *Variovorax*, and positive correlations with genera including *Ewingella, Stenotrophomonas*, and *Bacillus* (Table S6).

Because *Borrelia*-dominant samples were present within the tick-weak group, (5 of 19 samples), we examined whether *Borrelia*-specific correlation structure was driven by dominance events. To assess robustness, we repeated the analysis excluding *Borrelia*-dominant samples. Excluding these samples in fact increased overall network connectivity (62 nodes, 694 edges; density = 0.37; mean degree = 22.4 Fig. S4), and Borrelia associations persisted, indicating that the observed correlation structure was not driven solely by the dominance events (Fig. S4 and Table S7).

## Discussion

### Structure and diversity of the Ixodes scapularis microbiome

We have generated and analyzed full-length 16S rRNA gene sequences from 88 ticks sampled between 2022 and 2024 in Nova Scotia, Canada. The resulting microbiomes provide a wealth of information on the presence and abundance of the Lyme disease-causing bacterium *Borrelia burgdorferi* (present in 69% of ticks) relative to two other tick-adapted pathogens (*Anaplasma* (8.0%) and *Ehrlichia* (3.4%)) and environmentally acquired bacteria.

Alpha diversity patterns in the tick microbiomes were primarily driven by dominance intensity rather than the ecological identity of the dominant bacterial taxon. Shannon diversity and Pielou’s evenness are calculated directly from proportional abundances; therefore, strong dominance necessarily reduces entropy and constrains evenness independent of taxon origin. Samples classified as strongly dominated exhibited markedly reduced Shannon diversity and evenness, reflecting the mathematical properties of relative abundance distributions rather than ecological source per se. This distinction is important, as it prevents overinterpretation of alpha diversity differences as evidence of functional divergence between tick-adapted and environmentally dominated communities.

In contrast, beta diversity revealed a more complex structure. The ecological identity of the dominant taxon explained a larger proportion of community variation than dominance strength alone, and ordination analyses demonstrated that tick-adapted and environmentally dominated communities occupy distinct regions of compositional space. Dominance strength was found to modulate dispersion within these groups, with strongly dominated samples forming tighter clusters and weakly dominated samples exhibiting greater heterogeneity. Thus, microbiome organization in *Ixodes scapularis* appears structured along two partially independent axes, ecological source of dominance and dominance intensity.

### Environmental dominance as a structured community state

Environmentally dominated samples exhibited substantial internal co-occurrence structure, with network density and mean degree comparable to weakly dominated tick microbiome communities and far exceeding that observed in strongly tick-adapted samples. Importantly, this coordinated covariance among environmental taxa contrasts with the sparse networks characteristic of strongly dominated communities and is inconsistent with a purely stochastic contamination model.

Previous studies have often interpreted environmental bacterial taxa detected in tick microbiomes as transient background signal (Ross *et al*. 2018; Lejal *et al*. 2021). However, the coordinated positive and negative correlations detected herein among multiple environmental genera suggest structured community configurations rather than incidental presence/absence. While correlation-based networks cannot establish direct biological interaction, reproducible covariance patterns imply shared ecological drivers, such as exposure gradients, habitat-specific acquisition, or co-transmission from environmental reservoirs (Faust & Raes, 2012; Weiss *et al*. 2016).

It is important to note that bacterial taxa classified here as environmentally derived include those predicted to have been acquired from diverse external sources, including soil, vegetation, and vertebrate hosts. For example, genera such as *Staphylococcus* and *Streptococcus* are commonly associated with host skin and mucosa and may be introduced during blood feeding. However, the presence of coordinated covariance patterns among multiple taxa, including well-established soil- and plant-associated genera such as *Variovorax, Sphingomonas*, and *Methylobacterium*, supports the interpretation that environmentally dominated microbiomes reflect structured ecological assemblages rather than stochastic background signal.

Given that environmentally derived taxa may originate from multiple sources, one plausible driver of such structured environmental states is seasonal variation in tick behavior and exposure. Questing activity is known to vary across seasons and is associated with prolonged contact with soil and vegetation (Randolph *et al*. 2002; Eisen & Eisen 2018; Ogden *et al*. 2014). Coordinated environmental assemblages may therefore reflect ecological exposure regimes rather than random acquisition. Ongoing seasonal sampling and paired soil microbiome profiling will allow direct testing of whether environmental dominance and associated covariance structure track seasonal shifts in environmental microbial pools.

### *Borrelia* dominance and pathogen embedding

Although *Borrelia* is a tick-adapted pathogen, we found that tick microbiomes dominated by *Borrelia* clustered closer to environmentally dominated communities than to endosymbiont-dominated states in ordination space. This suggests that pathogen dominance represents a distinct and comparatively variable microbiome configuration relative to vertically transmitted endosymbiont dominance. Endosymbionts such as *Rickettsia* are intracellular and vertically transmitted, and often exhibit stable, constrained community structure (Sassera *et al*. 2006; Kurtti *et al*. 2015). In contrast, horizontally acquired pathogens may not impose the same structural constraints.

Vertically transmitted *Rickettsia* endosymbionts have been shown to frequently dominate the microbiome of *I. scapularis*—particularly in adult females—often comprising the vast majority of 16S rDNA reads and coinciding with reduced overall diversity (Thapa *et. al*. 2019a; Thapa *et. al*. 2019b; Van Treuren 2015; Gil *et. al*. 2020). Such extreme dominance can impose a compositional constraint that limits the detectable covariance structure among other taxa. In addition, experimental work suggests that *R. buchneri* may possess antibacterial or exclusion-like capabilities: tick cells hosting *R. buchneri* show reduced infectivity and replication for multiple tick-borne pathogens, consistent with the possibility that symbiont presence can constrain colonization by other bacteria (Cull *et al*. 2022).

Consistent with a dominance-constrained regime, we observed no *Borrelia* correlations exceeding |r| ≥ 0.3, in tick-strong communities. This pattern is expected when one taxon dominates the proportional abundance distribution (limiting variation in other taxa) and may be further reinforced if symbiont-associated exclusion reduces opportunities for stable co-variation. While our data do not demonstrate direct antagonism between *Rickettsia* and *Borrelia*, they support a model in which vertically transmitted symbiont dominance restricts the emergence of structured pathogen-associated covariance observed in weaker, more even community states (Narasimhan *et al*. 2014).

Co-occurrence analyses further indicated that pathogen embedding varies with microbiome community configuration. In weakly tick-dominated and environmentally dominated communities, *Borrelia* displayed multiple positive and negative associations with subsets of taxa. In environmentally dominated samples, *Borrelia* showed negative correlations with soil-associated genera such as *Variovorax, Caballeronia*, and *Luteibacter*, and positive correlations with genera including Erwinia and Legionella. In tick-weak samples, similar structured covariance was observed, including negative correlations with *Anaerococcus* and *Variovorax* and positive correlations with *Ewingella, Stenotrophomonas*, and *Bacillus*. Importantly, excluding *Borrelia*-dominant samples increased overall network connectivity and preserved correlation structure, indicating that these associations were not artifacts of *Borrelia*-dominance events. Rather, *Borrelia*-associated covariance emerges in contexts where community evenness permits coordinated variation among taxa.

These findings are consistent with prior work demonstrating that microbial community context can influence pathogen persistence and transmission potential in arthropod vectors (Narasimhan & Fikrig 2015; Abraham *et al*. 2017; Adegoke *et al*. 2020). While correlation does not imply direct interaction, state-dependent covariance patterns suggest that Borrelia embedding within the microbiome is contingent on broader community structure.

### Integrative model of microbiome states in Ixodes scapularis

Together, our results suggest that *I. scapularis* microbiomes oscillate between dominance-constrained states and coordinated multi-taxon assemblage states. Strongly dominated communities, particularly those dominated by vertically transmitted endosymbionts, exhibit reduced internal covariance and tight compositional clustering. In contrast, weakly tick-dominated and environmentally dominated communities display higher connectivity and coordinated structure.

Importantly, environmental dominance does not appear to represent stochastic background alone. Instead, the presence of structured covariance and reproducible network architecture suggests that environmentally dominated samples constitute a recurrent community configuration. Whether these states influence pathogen acquisition or persistence remains to be determined, but their structured nature argues against dismissing environmental taxa as mere contaminants.

One plausible ecological driver of these alternative dominance states is seasonal variation in tick behavior and exposure. Questing activity varies across the year and involves prolonged contact with soil and vegetation, potentially influencing both environmental microbial acquisition and the balance between tick-adapted and environmentally acquired taxa. Ongoing work sampling ticks across fall and spring, together with paired soil microbiome profiling from collection sites, will enable direct testing of whether environmental dominance and associated co-occurrence structure track seasonal changes in exposure and environmental microbial pools.

## Acknowledgements

The Archibald Lab acknowledges funding from a Genome Canada Emerging Issues Program grant (Lyme Disease in NS: The influence of strain variation on disease) led by Todd Hatchette, Nicholas Ogden, Robbin Lindsay, and Elizabeth Stringer. The authors appreciate the support of Archibald Lab members Ronie Haro, Dudley Chung, Dmytro Tymoshenko and Marlena Dlutek for assistance with tick sampling and laboratory workflows, and Laura Ferguson of Acadia University for advice on tick sampling.

