## Supplemental tables for "Full-length 16S ribosomal RNA gene sequencing reveals dynamics of tick-adapted and environmentally derived bacteria in the microbiome of the black-legged tick, *Ixodes scapularis* in Nova Scotia, Canada"

Table S1 DNA concentration in ng/µl of the 88 *Ixodes scapularis* tick samples.

| Samples | Concentration (ng/µl) |
|---|---|
| **adamo_001** | 6.92 |
| **adamo_002** | 4.51 |
| **adamo_003** | 5.51 |
| **adamo_004** | 3.44 |
| **adamo_005** | 3.39 |
| **adamo_006** | 3.63 |
| **adamo_007** | 6.88 |
| **adamo_008** | 8.57 |
| **adamo_009** | 6.77 |
| **adamo_010** | 8.18 |
| **adamo_011** | 4.44 |
| **adamo_012** | 4.9 |
| **adamo_013** | 3.9 |
| **adamo_014** | 5.46 |
| **adamo_015** | 4.94 |
| **adamo_016** | 3.46 |
| **adamo_017** | 4.92 |
| **adamo_018** | 8.76 |
| **adamo_019** | 3.92 |
| **adamo_020** | 5.7 |
| **adamo_021** | 4.28 |
| **adamo_022** | 7.54 |
| **adamo_023** | 4.92 |
| **adamo_024** | 5.8 |
| **adamo_025** | 4.92 |
| **adamo_026** | 8.26 |
| **adamo_027** | 6.56 |
| **adamo_028** | 4.24 |
| **adamo_029** | 5.06 |
| **adamo_030** | 3.8 |
| **adamo_031** | 5.08 |
| **adamo_032** | 4.54 |
| **adamo_033** | 4.26 |
| **adamo_034** | 5.54 |
| **adamo_035** | 5.06 |
| **adamo_036** | 2.86 |
| **adamo_037** | 3.78 |
| **adamo_038** | 3.94 |
| **adamo_039** | 2.14 |
| **adamo_040** | 4.56 |
| **adamo_041** | 2.64 |

| | |
|---|---|
| **adamo_042** | 3.14 |
| **adamo_043** | 2.74 |
| **adamo_044** | 3.16 |
| **adamo_045** | 3.84 |
| **adamo_046** | 3.46 |
| **adamo_047** | 4.14 |
| **adamo_048** | 3.06 |
| **adamo_049** | 3.02 |
| **adamo_050** | 2.98 |
| **adamo_051** | 1.742 |
| **adamo_052** | 1.49 |
| **adamo_053** | 2.46 |
| **adamo_054** | 6.48 |
| **adamo_055** | 4.42 |
| **adamo_056** | 2.74 |
| **adamo_057** | 3.62 |
| **adamo_058** | 3.6 |
| **adamo_059** | 2.88 |
| **adamo_060** | 5.32 |
| **adamo_061** | 3.72 |
| **adamo_062** | 3.1 |
| **adamo_063** | 3.14 |
| **adamo_064** | 2.28 |
| **adamo_065** | 14.18 |
| **adamo_066** | 12.4 |
| **adamo_067** | 7.6 |
| **adamo_068** | 8.76 |
| **adamo_069** | 16.06 |
| **adamo_070** | 8.28 |
| **adamo_071** | 9.9 |
| **adamo_072** | 10.42 |
| **adamo_073** | 9.66 |
| **adamo_074** | 11.16 |
| **adamo_075** | 9.48 |
| **adamo_076** | 13.22 |
| **adamo_077** | 14.76 |
| **adamo_078** | 13.26 |
| **adamo_079** | 9.54 |
| **adamo_080** | 12.72 |
| **adamo_081** | 10 |
| **archibald_001*** | 4.8 |
| **archibald_002*** | 14.2 |
| **archibald_003** | 5.91 |
| **archibald_004** | 11 |

| | |
|---|---|
| **archibald_005** | 16 |
| **archibald_006** | 12.2 |
| **archibald_007** | 18.3 |

*Male tick

Table S2 Numbers of reads pre- and post-quality control, genera, and alpha diversity (Shannon diversity, Pielou's evenness) measures for 88 *Ixodes scapularis* tick samples.

| **Samples** | **Raw Reads** | **Read Counts post QC** | **Number of genera** | **Shannon diversity** | **Pielou's evenness** |
|---|---|---|---|---|---|
| **adamo_001** | 241346 | 135420 | 63 | 2.92618564 | 0.65029098 |
| **adamo_002** | 609808 | 354261 | 19 | 0.15498001 | 0.05090454 |
| **adamo_003** | 479337 | 272864 | 31 | 1.09210929 | 0.3047589 |
| **adamo_004** | 167660 | 94058 | 73 | 2.35149372 | 0.51757454 |
| **adamo_005** | 22744 | 9549 | 41 | 2.38357045 | 0.62987606 |
| **adamo_006** | 137489 | 64475 | 58 | 2.36386345 | 0.55829097 |
| **adamo_007** | 106971 | 60090 | 33 | 1.54137943 | 0.42373689 |
| **adamo_008** | 293314 | 159086 | 27 | 0.94604556 | 0.28390981 |
| **adamo_009** | 86782 | 50927 | 40 | 1.82731319 | 0.47727457 |
| **adamo_010** | 43215 | 24403 | 16 | 0.48350349 | 0.17438702 |
| **adamo_011** | 603146 | 356432 | 39 | 2.07955555 | 0.55637757 |
| **adamo_012** | 241505 | 135791 | 34 | 1.57196578 | 0.4386654 |
| **adamo_013** | 675859 | 405830 | 33 | 0.91466231 | 0.25726379 |
| **adamo_014** | 44289 | 23962 | 19 | 1.43537912 | 0.48748815 |
| **adamo_015** | 684398 | 396375 | 20 | 0.19452949 | 0.0629333 |
| **adamo_016** | 1093273 | 684962 | 12 | 0.6671505 | 0.26848111 |
| **adamo_017** | 15538 | 9430 | 26 | 2.62025107 | 0.76303461 |
| **adamo_018** | 44082 | 15218 | 29 | 1.96165745 | 0.56103338 |
| **adamo_019** | 77253 | 44385 | 31 | 0.4374739 | 0.12622828 |
| **adamo_020** | 648026 | 372703 | 8 | 0.0352678 | 0.01696023 |
| **adamo_021** | 186604 | 127340 | 33 | 2.50824809 | 0.69993996 |
| **adamo_022** | 106026 | 72165 | 33 | 1.44815518 | 0.39528615 |
| **adamo_023** | 121031 | 84325 | 44 | 2.30164925 | 0.59780806 |
| **adamo_024** | 1035786 | 740871 | 19 | 0.84771676 | 0.27843998 |
| **adamo_025** | 374144 | 263884 | 25 | 0.43538321 | 0.1352594 |
| **adamo_026** | 276435 | 195125 | 20 | 0.72886945 | 0.2324576 |
| **adamo_027** | 432525 | 305530 | 33 | 1.22038958 | 0.3432546 |
| **adamo_028** | 459959 | 321025 | 21 | 0.40908292 | 0.12708876 |
| **adamo_029** | 1021409 | 725901 | 18 | 0.23087386 | 0.07841014 |
| **adamo_030** | 583027 | 416165 | 21 | 0.3550552 | 0.11486584 |
| **adamo_031** | 558444 | 396265 | 22 | 1.52661457 | 0.47426948 |
| **adamo_032** | 923282 | 646744 | 37 | 1.67790511 | 0.46126883 |
| **adamo_033** | 875687 | 622612 | 24 | 0.69693008 | 0.21390713 |
| **adamo_034** | 274149 | 198114 | 28 | 1.39604764 | 0.41459014 |
| **adamo_035** | 672837 | 468970 | 17 | 0.24317864 | 0.07987415 |
| **adamo_036** | 408436 | 326847 | 36 | 1.56127404 | 0.43568182 |
| **adamo_037** | 256071 | 197443 | 32 | 1.54798497 | 0.44272319 |
| **adamo_038** | 21200 | 15691 | 35 | 2.25411874 | 0.61105785 |
| **adamo_039** | 717291 | 558119 | 23 | 0.2388659 | 0.07420786 |

| | | | | | |
|---|---|---|---|---|---|
| **adamo_040** | 350934 | 276230 | 27 | 0.83435138 | 0.25315312 |
| **adamo_041** | 203412 | 152244 | 63 | 2.2474925 | 0.51896349 |
| **adamo_042** | 87721 | 67848 | 40 | 1.69567194 | 0.45083268 |
| **adamo_043** | 586825 | 454272 | 27 | 0.27627315 | 0.08045259 |
| **adamo_044** | 638857 | 498563 | 30 | 0.29008136 | 0.08226083 |
| **adamo_045** | 638598 | 490928 | 46 | 1.39275294 | 0.36174014 |
| **adamo_046** | 153173 | 119104 | 21 | 0.82500866 | 0.25959556 |
| **adamo_047** | 817314 | 636294 | 18 | 0.54796874 | 0.18610294 |
| **adamo_048** | 247794 | 191153 | 24 | 0.63638221 | 0.19532331 |
| **adamo_049** | 419302 | 319067 | 33 | 1.02385137 | 0.2814645 |
| **adamo_050** | 525641 | 410237 | 22 | 0.68132121 | 0.22041794 |
| **adamo_051** | 668652 | 502286 | 18 | 0.88207005 | 0.29444222 |
| **adamo_052** | 40886 | 31103 | 11 | 0.36661926 | 0.15289211 |
| **adamo_053** | 556961 | 421215 | 37 | 1.38935878 | 0.38476609 |
| **adamo_054** | 730159 | 547805 | 25 | 0.47627704 | 0.14450868 |
| **adamo_055** | 293223 | 222175 | 29 | 1.82736378 | 0.53727072 |
| **adamo_056** | 474222 | 344552 | 33 | 1.61424888 | 0.45403399 |
| **adamo_057** | 382047 | 292184 | 24 | 0.80935465 | 0.24556878 |
| **adamo_058** | 131317 | 99808 | 21 | 1.21712337 | 0.38817592 |
| **adamo_059** | 48619 | 36514 | 30 | 1.36084149 | 0.38590538 |
| **adamo_060** | 78634 | 59442 | 20 | 1.64399138 | 0.53185662 |
| **adamo_061** | 132744 | 100609 | 28 | 0.60038404 | 0.18017623 |
| **adamo_062** | 163189 | 126119 | 24 | 0.94087819 | 0.28878156 |
| **adamo_063** | 1221584 | 923116 | 3 | 0.00785104 | 0.00714633 |
| **adamo_064** | 215404 | 164183 | 19 | 1.03590802 | 0.34025304 |
| **adamo_065** | 1302872 | 991474 | 6 | 0.05002069 | 0.02791708 |
| **adamo_066** | 516841 | 385366 | 21 | 0.18174605 | 0.05879766 |
| **adamo_067** | 343294 | 255152 | 17 | 0.23308293 | 0.08064116 |
| **adamo_068** | 442560 | 333905 | 12 | 0.11727852 | 0.04719635 |
| **adamo_069** | 511927 | 382262 | 11 | 0.34224582 | 0.13772985 |
| **adamo_070** | 231612 | 171211 | 11 | 0.10986734 | 0.04421387 |
| **adamo_071** | 525983 | 394565 | 14 | 0.06596476 | 0.02499558 |
| **adamo_072** | 377883 | 283474 | 3 | 0.00250206 | 0.00227747 |
| **adamo_073** | 382665 | 283818 | 11 | 0.08941685 | 0.03598399 |
| **adamo_074** | 406078 | 306681 | 10 | 0.06222358 | 0.02702336 |
| **adamo_075** | 413760 | 312893 | 14 | 0.22169836 | 0.08400665 |
| **adamo_076** | 448839 | 335901 | 12 | 0.28478303 | 0.11460512 |
| **adamo_077** | 405945 | 295103 | 17 | 0.33138163 | 0.11696317 |
| **adamo_078** | 430568 | 324739 | 11 | 0.19324211 | 0.08058822 |
| **adamo_079** | 524612 | 394145 | 11 | 0.21337179 | 0.08898295 |
| **adamo_080** | 473662 | 351780 | 15 | 0.22900107 | 0.08456308 |
| **adamo_081** | 431054 | 324033 | 5 | 0.0327565 | 0.02035276 |
| **archibald_001*** | 678129 | 409640 | 38 | 1.62560055 | 0.41344675 |
| **archibald_002*** | 92225 | 57546 | 27 | 1.92333975 | 0.57118229 |

| | | | | | |
|---|---|---|---|---|---|
| **archibald_003** | 2186246 | 1560753 | 11 | 0.14695807 | 0.05914028 |
| **archibald_004** | 1634567 | 1220722 | 16 | 0.29100792 | 0.10495892 |
| **archibald_005** | 1453050 | 1087998 | 19 | 0.70182822 | 0.2383572 |
| **archibald_006** | 1347051 | 1023036 | 11 | 0.55197702 | 0.23019229 |
| **archibald_007** | 1323068 | 955986 | 4 | 0.19756279 | 0.14251143 |

*Male tick

Table S3. Dominance analysis of 88 *Ixodes scapularis* tick samples.

| Samples | Berger-Parker index | Dominant taxon | Second taxon | Dominance margin | Community | Dominance strength (0.5 margin) |
|---|---|---|---|---|---|---|
| **adamo_001** | 0.1955 | *Rickettsia* | *Paraburkholderia* | 0.0445 | tick | weak |
| **adamo_002** | 0.9772 | *Rickettsia* | *Borrelia* | 0.9648 | tick | strong |
| **adamo_003** | 0.7716 | *Rickettsia* | *Borrelia* | 0.6923 | tick | strong |
| **adamo_004** | 0.3301 | *Methylobacterium* | *Sphingomonas* | 0.1275 | env | weak |
| **adamo_005** | 0.3291 | *Erwinia* | *Borrelia* | 0.1660 | env | weak |
| **adamo_006** | 0.4560 | *Rickettsia* | *Luteibacter* | 0.3571 | tick | weak |
| **adamo_007** | 0.4173 | *Erwinia* | *Caballeronia* | 0.0247 | env | weak |
| **adamo_008** | 0.7479 | *Rickettsia* | *Caballeronia* | 0.6099 | tick | strong |
| **adamo_009** | 0.3569 | *Pseudomonas* | *Luteibacter* | 0.0090 | env | weak |
| **adamo_010** | 0.9045 | *Rickettsia* | *Luteibacter* | 0.8590 | tick | strong |
| **adamo_011** | 0.3137 | *Pseudomonas* | *Caballeronia* | 0.1277 | env | weak |
| **adamo_012** | 0.4739 | *Rickettsia* | *Pseudomonas* | 0.2437 | tick | weak |
| **adamo_013** | 0.5433 | *Rickettsia* | *Pseudomonas* | 0.1226 | tick | weak |
| **adamo_014** | 0.6068 | *Borrelia* | *Caballeronia* | 0.4554 | tick | weak |
| **adamo_015** | 0.9696 | *Rickettsia* | *Pseudomonas* | 0.9527 | tick | strong |
| **adamo_016** | 0.7297 | *Pseudomonas* | *Erwinia* | 0.4784 | env | weak |
| **adamo_017** | 0.2125 | *Bacillus* | *Luteibacter* | 0.0857 | env | weak |
| **adamo_018** | 0.4239 | *Borrelia* | *Sinocapsa* | 0.2468 | tick | weak |
| **adamo_019** | 0.9326 | *Rickettsia* | *Caballeronia* | 0.9168 | tick | strong |
| **adamo_020** | 0.9956 | *Rickettsia* | *Borrelia* | 0.9940 | tick | strong |
| **adamo_021** | 0.3084 | *Borrelia* | *Caballeronia* | 0.1600 | tick | weak |
| **adamo_022** | 0.6651 | *Anaplasma* | *Caballeronia* | 0.5673 | tick | strong |
| **adamo_023** | 0.3979 | *Borrelia* | *Pseudomonas* | 0.2272 | tick | weak |
| **adamo_024** | 0.6818 | *Rickettsia* | *Pseudomonas* | 0.4091 | tick | weak |
| **adamo_025** | 0.9145 | *Rickettsia* | *Pseudomonas* | 0.8670 | tick | strong |
| **adamo_026** | 0.8421 | *Rickettsia* | *Borrelia* | 0.7868 | tick | strong |
| **adamo_027** | 0.6834 | *Rickettsia* | *Pseudomonas* | 0.5322 | tick | strong |
| **adamo_028** | 0.9303 | *Rickettsia* | *Pseudomonas* | 0.8916 | tick | strong |

| | | | | | | |
|---|---|---|---|---|---|---|
| **adamo_029** | 0.9623 | *Rickettsia* | *Pseudomonas* | 0.9451 | tick | strong |
| **adamo_030** | 0.9373 | *Rickettsia* | *Pseudomonas* | 0.9124 | tick | strong |
| **adamo_031** | 0.4629 | *Borrelia* | *Pseudomonas* | 0.1560 | tick | weak |
| **adamo_032** | 0.3772 | *Rickettsia* | *Pseudomonas* | 0.1037 | tick | weak |
| **adamo_033** | 0.8355 | *Rickettsia* | *Pseudomonas* | 0.7706 | tick | strong |
| **adamo_034** | 0.4139 | *Pseudomonas* | *Luteibacter* | 0.0132 | env | weak |
| **adamo_035** | 0.9559 | *Rickettsia* | *Pseudomonas* | 0.9317 | tick | strong |
| **adamo_036** | 0.3764 | *Ehrlichia* | *Anaplasma* | 0.0261 | tick | weak |
| **adamo_037** | 0.3569 | *Caballeronia* | *Pseudomonas* | 0.0211 | env | weak |
| **adamo_038** | 0.3523 | *Streptococcus* | *Lactococcus* | 0.1403 | env | weak |
| **adamo_039** | 0.9630 | *Rickettsia* | *Pseudomonas* | 0.9466 | tick | strong |
| **adamo_040** | 0.7888 | *Rickettsia* | *Erwinia* | 0.6836 | tick | strong |
| **adamo_041** | 0.5620 | *Caballeronia* | *Paraburkholderia* | 0.5145 | env | strong |
| **adamo_042** | 0.6456 | *Rickettsia* | *Borrelia* | 0.5774 | tick | strong |
| **adamo_043** | 0.9599 | *Rickettsia* | *Staphylococcus* | 0.9502 | tick | strong |
| **adamo_044** | 0.9590 | *Rickettsia* | *Borrelia* | 0.9488 | tick | strong |
| **adamo_045** | 0.6267 | *Rickettsia* | *Pseudomonas* | 0.4684 | tick | weak |
| **adamo_046** | 0.8324 | *Pseudomonas* | *Luteibacter* | 0.7529 | env | strong |
| **adamo_047** | 0.8579 | *Rickettsia* | *Pseudomonas* | 0.7465 | tick | strong |
| **adamo_048** | 0.8624 | *Rickettsia* | *Pseudomonas* | 0.7990 | tick | strong |
| **adamo_049** | 0.7403 | *Pseudomonas* | *Rickettsia* | 0.6355 | env | strong |
| **adamo_050** | 0.8365 | *Rickettsia* | *Pseudomonas* | 0.7425 | tick | strong |
| **adamo_051** | 0.7303 | *Pseudomonas* | *Luteibacter* | 0.5438 | env | strong |
| **adamo_052** | 0.9301 | *Rickettsia* | *Borrelia* | 0.9107 | tick | strong |
| **adamo_053** | 0.4386 | *Rickettsia* | *Caballeronia* | 0.0694 | tick | weak |
| **adamo_054** | 0.9170 | *Rickettsia* | *Pseudomonas* | 0.8940 | tick | strong |
| **adamo_055** | 0.4200 | *Rickettsia* | *Pseudomonas* | 0.2163 | tick | weak |
| **adamo_056** | 0.4686 | *Rickettsia* | *Pseudomonas* | 0.2772 | tick | weak |
| **adamo_057** | 0.6436 | *Rickettsia* | *Pseudomonas* | 0.3104 | tick | weak |
| **adamo_058** | 0.6054 | *Rickettsia* | *Pseudomonas* | 0.3762 | tick | weak |
| **adamo_059** | 0.6935 | *Rickettsia* | *Pseudomonas* | 0.6022 | tick | strong |

| | | | | | | |
|---|---|---|---|---|---|---|
| **adamo_060** | 0.4598 | *Pseudomonas* | *Rickettsia* | 0.2040 | env | weak |
| **adamo_061** | 0.8853 | *Rickettsia* | *Borrelia* | 0.8492 | tick | strong |
| **adamo_062** | 0.7277 | *Pseudomonas* | *Anaplasma* | 0.5347 | env | strong |
| **adamo_063** | 0.9991 | *Pseudomonas* | *Rickettsia* | 0.9983 | env | strong |
| **adamo_064** | 0.6036 | *Rickettsia* | *Pseudomonas* | 0.2882 | tick | weak |
| **adamo_065** | 0.9930 | *Rickettsia* | *Pseudomonas* | 0.9890 | tick | strong |
| **adamo_066** | 0.9702 | *Rickettsia* | *Pseudomonas* | 0.9520 | tick | strong |
| **adamo_067** | 0.9580 | *Rickettsia* | *Pseudomonas* | 0.9302 | tick | strong |
| **adamo_068** | 0.9820 | *Rickettsia* | *Pseudomonas* | 0.9715 | tick | strong |
| **adamo_069** | 0.9190 | *Rickettsia* | *Pseudomonas* | 0.8501 | tick | strong |
| **adamo_070** | 0.9933 | *Rickettsia* | *Erwinia* | 0.9916 | tick | strong |
| **adamo_071** | 0.9917 | *Rickettsia* | *Pseudomonas* | 0.9891 | tick | strong |
| **adamo_072** | 0.9997 | *Rickettsia* | *Neochroococcus* | 0.9996 | tick | strong |
| **adamo_073** | 0.9855 | *Rickettsia* | *Pseudomonas* | 0.9738 | tick | strong |
| **adamo_074** | 0.9908 | *Rickettsia* | *Caballeronia* | 0.9842 | tick | strong |
| **adamo_075** | 0.9575 | *Rickettsia* | *Pseudomonas* | 0.9261 | tick | strong |
| **adamo_076** | 0.9378 | *Rickettsia* | *Pseudomonas* | 0.8879 | tick | strong |
| **adamo_077** | 0.9354 | *Rickettsia* | *Pseudomonas* | 0.8978 | tick | strong |
| **adamo_078** | 0.9653 | *Rickettsia* | *Pseudomonas* | 0.9434 | tick | strong |
| **adamo_079** | 0.9578 | *Rickettsia* | *Pseudomonas* | 0.9251 | tick | strong |
| **adamo_080** | 0.9520 | *Rickettsia* | *Pseudomonas* | 0.9117 | tick | strong |
| **adamo_081** | 0.9952 | *Rickettsia* | *Pseudomonas* | 0.9910 | tick | strong |
| **archibald_001*** | 0.5294 | *Caballeronia* | *Sphingomonas* | 0.3371 | env | weak |
| **archibald_002*** | 0.3633 | *Sphingomonas* | *Caballeronia* | 0.0450 | env | weak |
| **archibald_003** | 0.9724 | *Rickettsia* | *Borrelia* | 0.9491 | tick | strong |
| **archibald_004** | 0.9458 | *Rickettsia* | *Anaplasma* | 0.9220 | tick | strong |
| **archibald_005** | 0.7840 | *Rickettsia* | *Caballeronia* | 0.6162 | tick | strong |
| **archibald_006** | 0.8063 | *Rickettsia* | *Pseudomonas* | 0.6236 | tick | strong |
| **archibald_007** | 0.9523 | *Pseudomonas* | *Rickettsia* | 0.9058 | env | strong |

*Male tick

Table S4. Kruskal-Wallis rank sum test of alpha diversity measures compared between community source (tick-adapted vs environmentally acquired), dominance strength (strong vs weak) at both margin 0.5 and 0.3, and community and dominance strength (env-weak, env-strong, tick-weak, tick-strong) at both 0.5 and 0.3 margin. Significant (<0.05) pairwise results are shown in parentheses in order of significance.

| | **Shannon diversity** | **Pielou's evenness** |
|---|---|---|
| **community** | $\chi^2 = 12.6$, df = 1, $p = 3.78 \times 10^{-4}$ | $\chi^2 = 14.2$, df = 1, $p = 1.61 \times 10^{-4}$ |
| **dominance (0.5)** | $\chi^2 = 48.52$, df = 1, $p = 3.28 \times 10^{-12}$ | $\chi^2 = 50.47$, df = 1, $p = 1.21 \times 10^{-12}$ |
| **dominance (0.3)** | $\chi^2 = 41.43$, df = 1, $p = 1.22 \times 10^{-10}$ | $\chi^2 = 42.4$, df = 1, $p = 7.44 \times 10^{-11}$ |
| **community – dominance (0.5)** | $\chi^2 = 50.88$, df = 3, $p = 5.20 \times 10^{-11}$ (ts-tw, ts-ew, es-ew, es-tw) | $\chi^2 = 53.71$, df = 3, $p = 1.29 \times 10^{-11}$ (ts-tw, ts-ew, es-ew, es-tw, ts-es) |
| **community – dominance (0.3)** | $\chi^2 = 44.01$, df = 3, $p = 1.5 \times 10^{-09}$ (ts-tw, ts-ew, es-ew, es-tw, ts-es) | $\chi^2 = 45.82$, df = 3, $p = 6.20 \times 10^{-10}$ (ts-tw, ts-ew, es-ew, es-tw, ts-es) |

ts tick-strong; tw tick-weak; es environment-strong; ew environment-weak

Table S5. Taxa correlated with *Borrelia* (|r| ≥ 0.3) in tick samples dominated by environmentally acquired taxa.

| Taxa | Environmental (r) |
|---|---|
| *Mycobacterium* | -0.44 |
| *Variovorax* | -0.42 |
| *Caballeronia* | -0.38 |
| *Luteibacter* | -0.38 |
| *Cutibacterium* | -0.37 |
| *Lichenifustis* | -0.36 |
| *Rhizobacter* | -0.35 |
| *Hymenobacter* | -0.33 |
| *Paludisphaera* | -0.30 |
| *Paraburkholderia* | 0.34 |
| *Rhodanobacter* | 0.35 |
| *unassigned* | 0.37 |
| *Erwinia* | 0.44 |
| *Legionella* | 0.59 |

Table S6. Taxa correlated with *Borrelia* (|r| ≥ 0.3) in tick samples weakly dominated by Tick-adapted taxa

| Taxa | Environmental (r) |
|---|---|
| *Anaerococcus* | -0.63 |
| *Variovorax* | -0.61 |
| *Schauerella* | -0.53 |
| *Methylobacterium* | -0.49 |
| *Terriglobus* | -0.39 |
| *Curtobacterium* | -0.36 |
| *Bradyrhizobium* | -0.34 |
| *Rhizobacter* | -0.32 |
| *Rickettsia* | -0.31 |
| *Caldimonas* | -0.31 |
| *unassigned* | 0.30 |
| *Paraburkholderia* | 0.31 |
| *Globicatella* | 0.32 |
| *Bacillus* | 0.35 |
| *Caballeronia* | 0.39 |
| *Leclercia* | 0.42 |
| *Stenotrophomonas* | 0.45 |
| *Ewingella* | 0.52 |

Table S7. Taxa correlated with *Borrelia* (|r| ≥ 0.3) in tick samples weakly dominated by Tick-adapted taxa, excluding tick samples where *Borrelia* was the dominate *taxa*.

| **Taxa** | **Environmental (r)** |
|---|---|
| *Anaerococcus* | -0.67 |
| *Variovorax* | -0.65 |
| *Methylobacterium* | -0.51 |
| *Comamonas* | -0.50 |
| *Rhizobacter* | -0.46 |
| *Bradyrhizobium* | -0.46 |
| *Schauerella* | -0.39 |
| *Methylocapsa* | -0.35 |
| *Lichenibacterium* | -0.34 |
| *Achromobacter* | -0.33 |
| *Frigoribacterium* | -0.32 |
| *Terriglobus* | -0.31 |
| *Curtobacterium* | -0.30 |
| *Phyllobacterium* | 0.32 |
| *Thermoleptolyngbya* | 0.32 |
| *Caballeronia* | 0.35 |
| *Pseudomonas* | 0.37 |
| *Ewingella* | 0.47 |
| *Stenotrophomonas* | 0.53 |
| *Bacillus* | 0.55 |
| *Leclercia* | 0.61 |
| *Erwinia* | 0.64 |
