## Supplemental figure legends for "Full-length 16S ribosomal RNA gene sequencing reveals dynamics of tick-adapted and environmentally derived bacteria in the microbiome of the black-legged tick, *Ixodes scapularis* in Nova Scotia, Canada"

**Figure S1. Alpha diversity plots by ecological category of the dominant taxa in each tick (tick-adapted or environmental) (A)** Shannon diversity ($\chi^2 = 12.6$, df = 1, p = $3.78 \times 10^{-4}$). **(B)** Pielou's evenness ($\chi^2 = 14.2$, df = 1, p = $1.61 \times 10^{-4}$). Dominance was defined by a marginal difference between top two most abundant taxa with strong $\geq 0.5$ and weak dominance $<0.5$. Colors of dots represent dominance strength with black being strong and grey weak.

**Figure S2. Alpha diversity plots by ecological category of the dominant taxa in each tick (tick-adapted or environmental) and strength of dominance (strong or weak). (A)** Shannon diversity ($\chi^2 = 50.88$, df = 3, p = $5.20 \times 10^{-11}$). **(B)** Pielou's evenness ($\chi^2 = 53.71$, df = 3, p = 1.29 $\times 10^{-11}$). Dominance was defined by a marginal difference between top two most abundant taxa with strong $\geq 0.5$ and weak dominance $<0.5$. Colors of dots represent ecological category and dominance strength, lime green = environmental-strong dominance; teal green=environmental-weak dominance; brown=tick-adapted-strong dominance; orange=tick-adapted-weak dominance.

**Figure S3. Principle coordinate analysis plot of Bray Curtis dissimilarities among 88 *I. scapularis* tick samples. (A)** Tick samples are colored based on the dominance structure as measured by the difference in abundance of the top two taxa. Black = strong dominance (difference $\geq 0.5$); grey= weak dominance (difference $< 0.5$). (**B)** Tick samples are colored based on category of assigned dominant taxa, brown=tick-adapted; green=environmental. In both plots PC1 and PC2 explained 52.6% and 16.6% of the variance.

**Figure S4. Co-occurrence networks for tick samples that were weakly dominated** (**margin <0.5) by tick-adapted taxa, but excluding those tick samples dominated by *Borrelia*.** Only taxa present in ≥20% of samples were retained, count data were transformed using centered log-ratio (CLR), and Pearson correlations were calculated. Only edges with a correlation threshold of $|r| \geq 0.3$ were retained. The network consisted of 62 nodes, 694 edges and had a density = 0.37, and mean degree = 22.4. Node size is proportional to within-network degree (normalized independently for each network). The node corresponding to *Borrelia* is colored orange with all other nodes colored grey. Edges corresponding to positive correlations are shown in blue, and negative correlations in red.
