## Supplementary figures and images for "Full-length 16S ribosomal RNA gene sequencing reveals dynamics of tick-adapted and environmentally derived bacteria in the microbiome of the black-legged tick, *Ixodes scapularis* in Nova Scotia, Canada"

### Supplemental figures

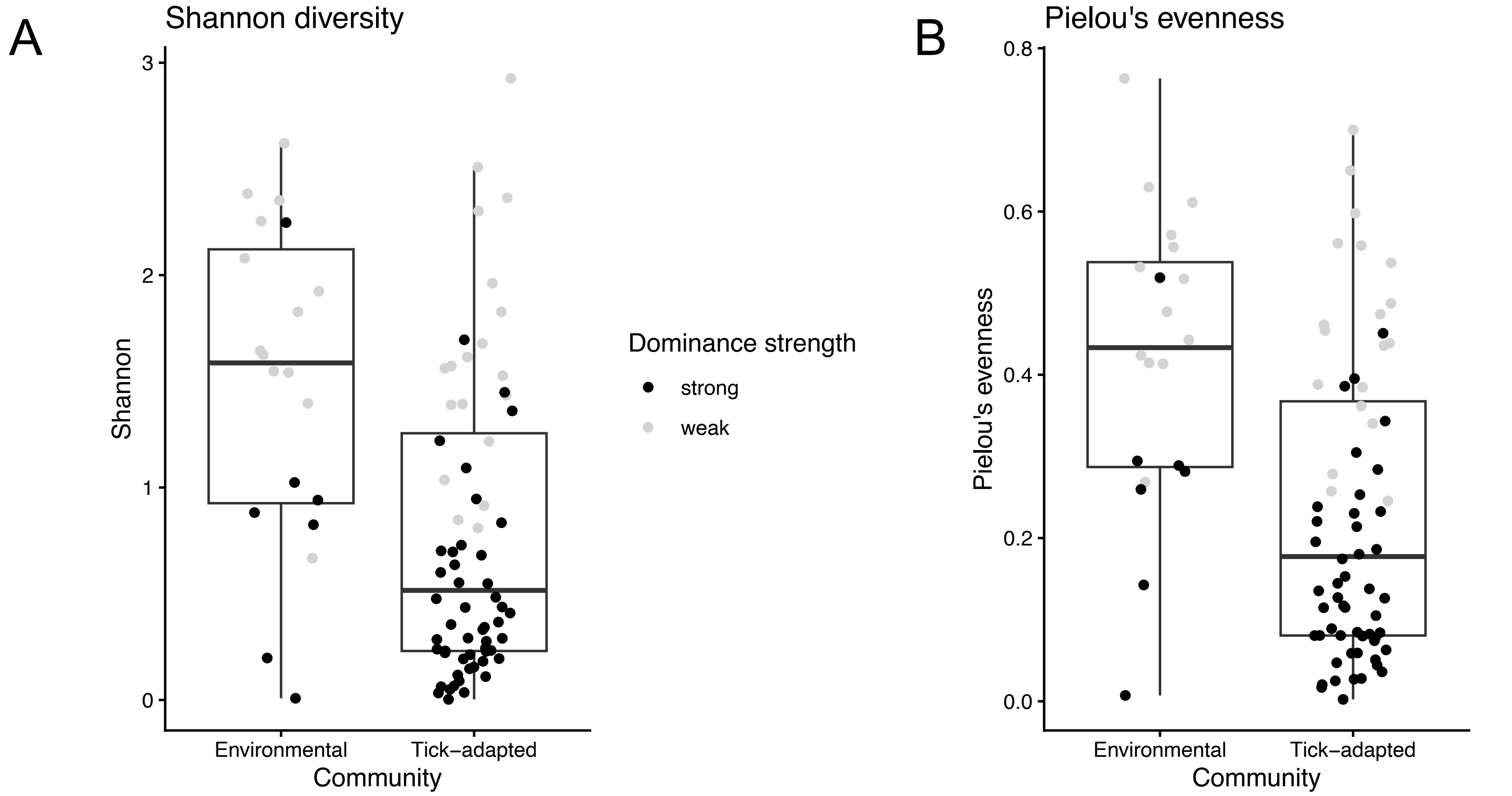

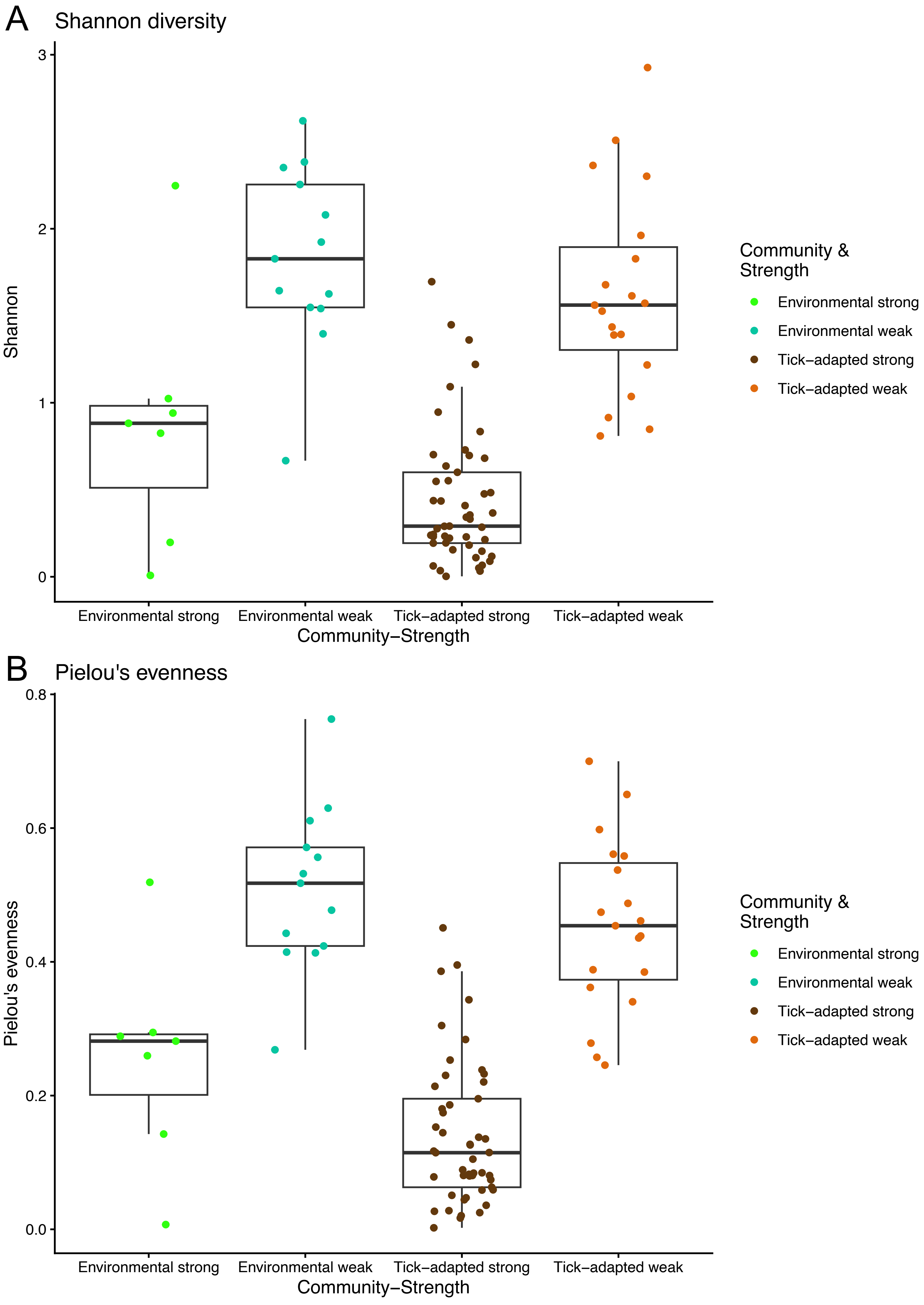

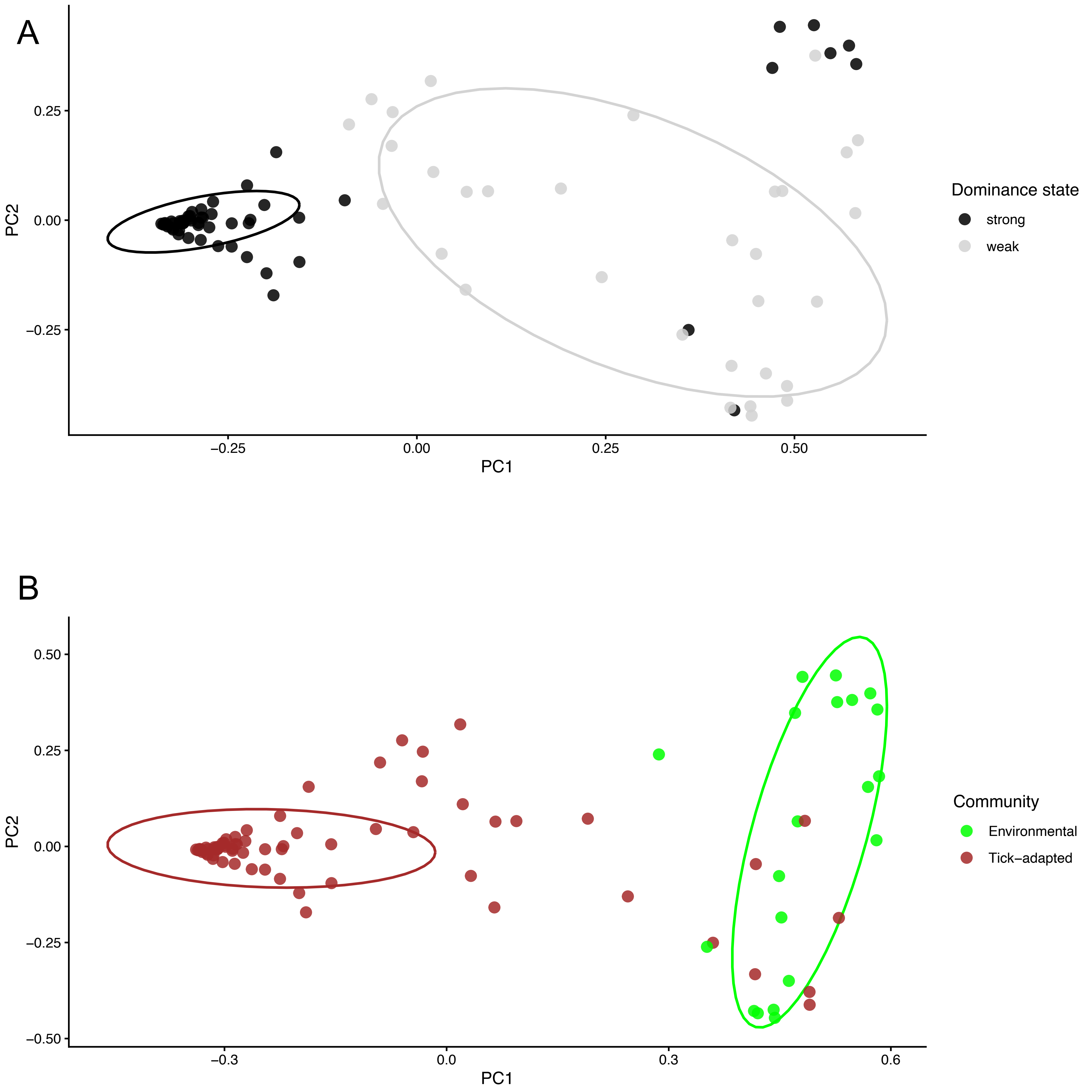

Tick–weak no *Borrelia*

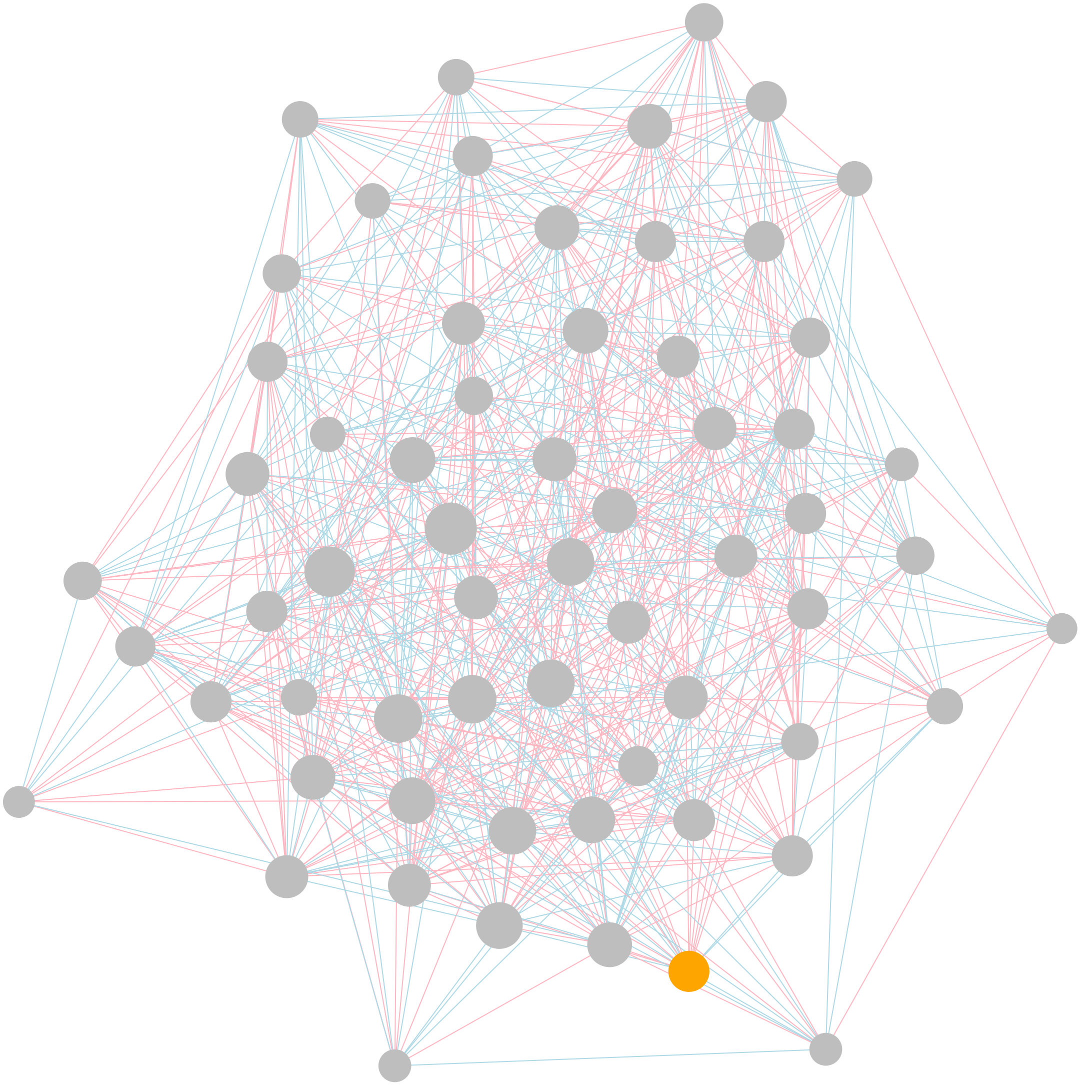
